# Structure of a type IV CRISPR-Cas effector complex

**DOI:** 10.1101/2020.07.31.231399

**Authors:** Yi Zhou, Jack P.K. Bravo, Hannah N. Taylor, Jurre Steens, Ryan N. Jackson, Raymond H.J. Staals, David W. Taylor

## Abstract

We reveal the structure of a type IV-B CRISPR effector (Csf) complex at 3.9 Å resolution using cryo-electron microscopy. The complex resembles the type III-A CRISPR Csm effector complex, but lacks subunits for RNA processing and target DNA cleavage, and is surprisingly assembled upon heterogeneous non-CRISPR RNA. These findings provide the first glimpse into the assembly and function of enigmatic type IV CRISPR systems.

---

Bacteria and archaea employ CRISPR (Clustered Regularly Interspaced Short Palindromic Repeat)-Cas (CRISPR-associated) systems for adaptive immunity against phages, plasmids and other mobile-genetic elements (MGEs)^1^. In the multi-subunit class I systems, the CRISPR locus is transcribed and processed into small crRNA guides (CRISPR-derived RNA), around which several Cas proteins assemble to form large ribonucleoprotein complexes that facilitate RNA-guided surveillance and degradation of complementary targets^2^. While a myriad of structures have been determined for most types of CRISPR RNA-guided complexes (types I^3–8^, II^9–11^, III^12–14^, V^15–17^, VI^18–20^), the RNA complexes of the highly diverse class 1 type IV CRISPR systems have largely remained structurally uncharacterised^1,21,22^. Type IV CRISPR systems occur within plasmid-like elements and lack genes encoding adaptation modules (Cas1, Cas2 and Cas4)^23,24^. The *Mycobacterium sp*. JS623 subtype IV-B CRISPR operon is encoded within a megaplasmid, and lacks both a pre-crRNA maturase (Cas6/Csf5^25^) and a CRISPR array, containing only *csf1* (large subunit), *csf4* (small subunit), *csf2* (Cas7) and *csf3* (Cas5) genes (**Figure 1a**). Interestingly, *M. sp. JS623* also harbours a type I-E system (with an associated CRISPR array) on the same megaplasmid, and another type IV-B operon encoded on a different megaplasmid (**Supplementary Figure 1**), suggesting that type IV-B complexes may assemble on other crRNAs. However, the structure and function of such hybrid complexes are unknown.

**Figure 1.**
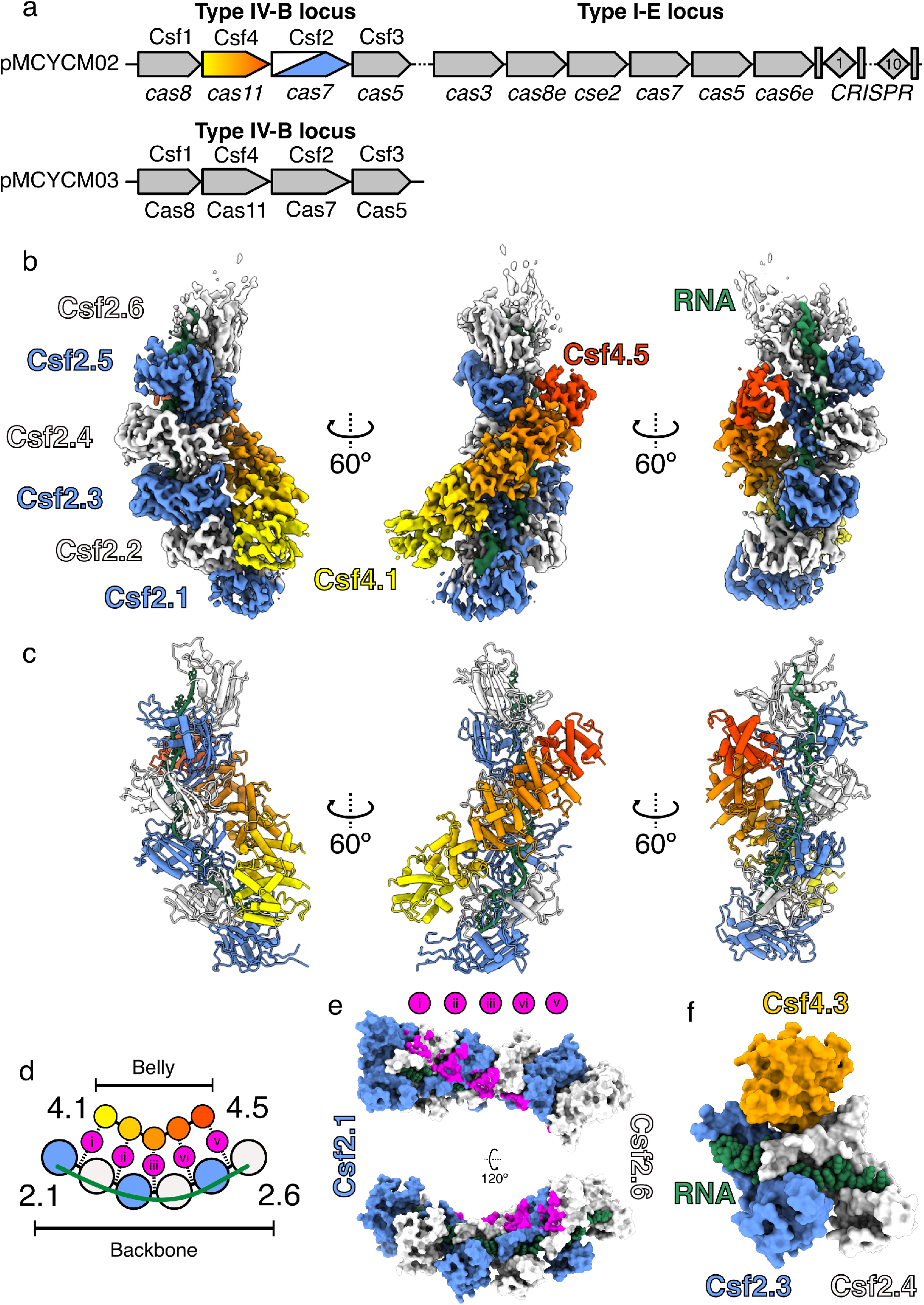
Structure of type IV-B CRISPR complex. **a**, *M. sp*. JS623 plasmid-encoded CRISPR operons. Top: Type IV-B and I-E CRISPR loci present on pMCYCM02 megaplasmid. Bottom: Additional type IV-B locus encoded by pMCYCM03 megaplasmid. **b**, 3.9 Å-resolution cryo-EM reconstruction of type IV-B CRISPR complex. Cas7 subunits are colored blue and white, and five Cas11 subunits are colored as a yellow-orange-red gradient. Csf-bound RNA is green. **c**, Refined model for the Csf effector complex derived from the cryo-EM maps shown in **b**. **d**, Schematic of Cas7-Cas11 interactions. Five Csf2-Csf4 interactions occur in this complex (labelled i – v). **e**, Positions of Csf4 contacts on Csf2 backbone, colored magenta as shown in panel. **d**, Csf4 sits upon the Csf2-Csf2 interface. **f**, Csf4 binds a the interface with buried surface area of 505 Å^2^ (150 Å^2^ and 355 Å^2^ with Csf2.3 and Csf2.4, respectively). Csf4 is completely occluded from bound RNA. Csf2 subunits are intimately connected (1021 Å^2^) and make a network of contacts with bound RNA (~1200 Å^2^ buried surface area per Csf2 subunit).

To gain mechanistic insights into the type IV-B system, we first transformed *E. coli* BL21 cells with an expression plasmid encoding the *M. sp*. JS623 type IV-B Cas proteins, and the *M. sp*. JS623 type I-E Cas6 and associated CRISPR array (**Figure 1a**). Using strep-tag affinity and size exclusion chromatography, we observed filamentous ribonucleoprotein (RNP) complexes that eluted close to the void volume^24^ and a smaller, discrete, RNA-containing species reminiscent of class 1 multi-subunit crRNA-guided complexes (**Supplementary figure 1**)^26^. While this latter fraction contained all four Csf subunits, Csf2 and Csf4 were the most abundant (**Supplementary figure 1**). Despite the appearance of a uniform band of ~55-60 nucleotides on denaturing PAGE (**Supplementary figure 2a**), RNAseq analysis revealed bound RNAs were heterogeneous in nature. Few RNAs were derived from the plasmid-encoded CRISPR array, while the majority of Csf-bound RNAs originated from the expression plasmid (63%). (**Supplementary figure 2b**). Surprisingly, classification of Csf-bound RNAs showed that most were derived from non-coding RNAs, with enrichment for small (regulatory) RNAs (sRNAs) (**Supplementary figure 2c; Supplementary File 1**)^27^. The apparent lack of sequence specific assembly of the Csf complex on a crRNA is different from other CRISPR-Cas systems^26^, and the enrichment of sRNA in place of a crRNA guide suggests IV-B complexes may have evolved a unique regulatory function.

To compare the type IV-B RNA-guided complex to the complexes of other class 1 systems we next determined a cryo-EM structure of the IV-B Csf complex at 3.9 Å resolution (**Figure 1b, Supplementary Figure 3**), allowing us to build an atomic model of the complex *de novo* (**Figure 1c**). The type IV-B complex resembles a sea cucumber, with seven Csf2 (Cas7-like) subunits forming a helical ‘backbone,’ while five Csf4 (Cas11) subunits comprise a helical ‘belly’. Each Csf4 subunit sits upon a Csf2-Csf2 interface (**Figure 1d, e & f**). The “helical bundle” topology of Csf4 (**Supplementary Figure 4**) and presence of a contiguous positively-charged patch running along the length of the minor filament (**Supplementary Figure 5**) are typical of Cas small subunits in class 1 CRISPR systems^3,12^.

Csf2 makes extensive contacts with the bound RNA (buried surface area of ~1200 Å^2^ per Csf2 subunit) via a positively-charged palm (**Figure 2a**). The thumb of the neighbouring Csf2 subunit protrudes into the centre of the palm, inducing a kink in the RNA backbone and a ‘flipped’ base at six nucleotide intervals (typical of other class 1 complexes^14^). Using our atomic model of Csf2 we searched for structural homologues. Csf2 had significant similarity to the type III-A CRISPR Csm3 (i.e. Cas7) subunit (Dali Z-score of 14.1), despite a sequence identity of only 16%. Csf2 and Csm3 superimpose with an r.m.s.d of 2.9 Å and use equivalent interfaces to bind RNA and induce near-identical RNA backbone conformations (r.m.s.d of 1.5 Å) (**Figure 2a**). This supports previous bioinformatics-based hypotheses that type IV systems originated from type III-like ancestors^1,23,24^.

**Figure 2.**
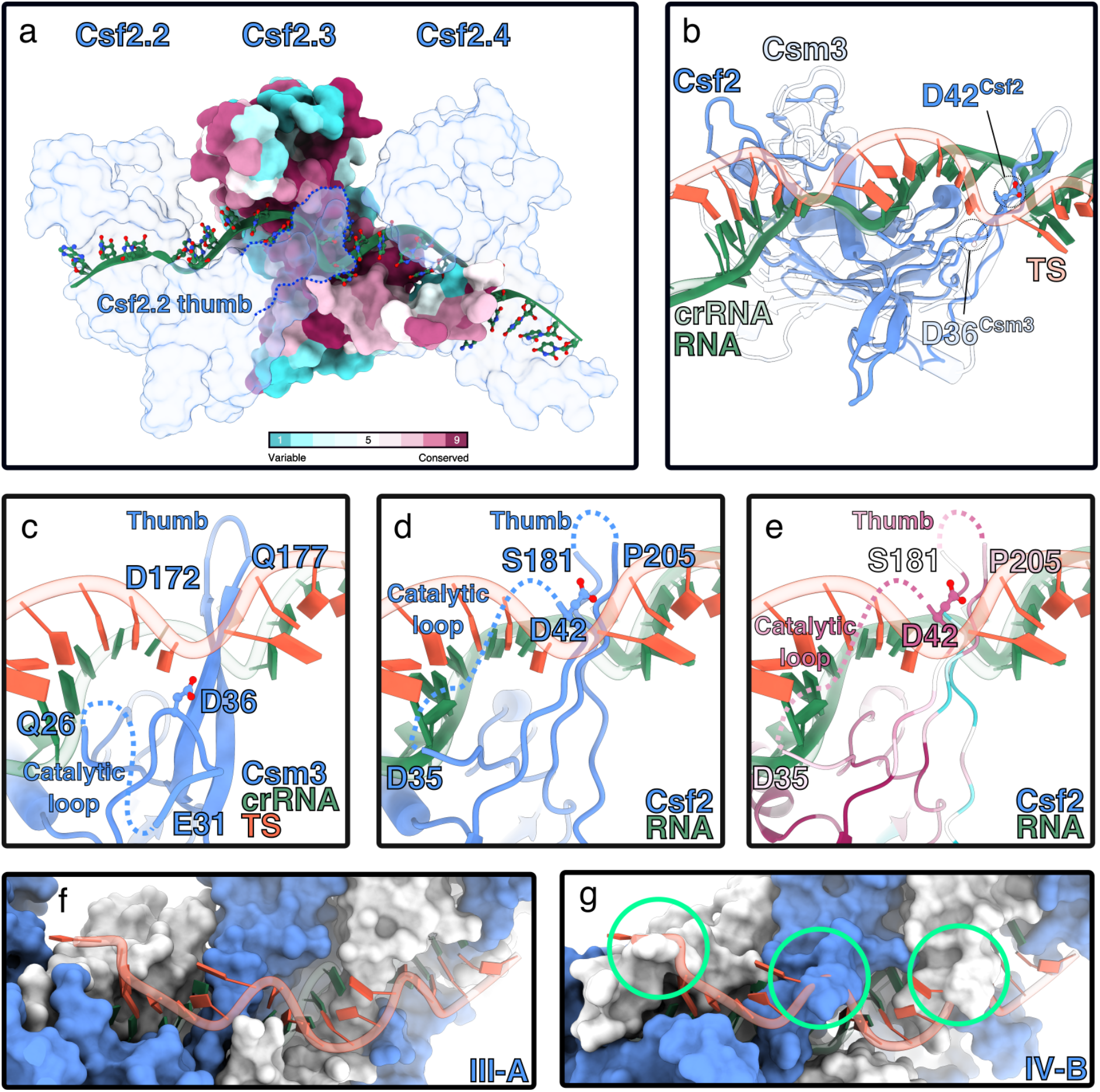
RNA-binding by type IV-B Cas7. **a**, RNA (green) binding site runs across the palms of Csf2 subunits. Csf2.3 is colored according to conservation. The “thumb” of the n-1 Csf2 (i.e. Csf2.2) protrudes into the backbone of bound RNA (solid green), inducing a kink. **b**, Alignment of type III-A backbone subunit Csm3 (PDB 607i, transparent) with Csf2 (solid blue). Csm3 and Csf2 align with an r.m.s.d. of 2.9Å, with a Dali server Dali server Z-score 14.1. Csf2-bound RNA binds in the same conformation as crRNA (transparent green) to Csm3 (RMSD of 1.5 Å). Catalytic residue Asp36^Csm3^ and putative catalytic residue Asp42^Csf2^ side chains are located near the target strand (TS - transparent red), bound to the type III crRNA (transparent green). **c**, Residues flanking unstructured catalytic loop (27-35) and apical loop of Csm3 thumb also interact with the TS. Catalytic residue D36 is shown for clarity. **d & e**, Putative interactions with Csf2 and TS, based on alignments with the Csm complex. e is colored by conservation. The Csf2 thumb contains a flexible 20 residue insertion, not visible in our cryo-EM map. **f & g**, Path of TS bound by type III-A Csm complex (**f**) and putative path along IV-B (**g**). Severe clashes with TS Csf2 are circled in green.

The Type III backbone protein Csm3 has RNase activity, and cleaves the phosphodiester backbone of the crRNA-bound target strand (TS) at 6-nt intervals^28^. Given that the Csm crRNA aligns almost perfectly with Csf-bound RNA, we reasoned that Csf2 might also possess RNase activity. Within our aligned structures, both the catalytic Asp36^Csm3^ residue and the conserved Asp42^Csf2^ residue are similarly positioned within an unstructured “catalytic loop” (**Figure 2c-e, Supplementary figure 6**). However, despite this similarity, in our structure Asp42^Csf2^ is not in a position compatible with target RNA cleavage. Additionally, further structural alignment reveals significant steric clashes between the path of a bound target sequence (TS) and the Csf2 catalytic loop and the inferred position of an unstructured 24-residue insertion within the Csf2 thumb (**Figure 2f & g**), suggesting a significant conformational rearrangement of subunits would need to occur to place D42^Csf2^ in a position amenable to catalyse target RNA cleavage.

Our structure of the Csf complex provides evidence that type IV-B evolved from type III CRISPR-Cas systems, but lost its CRISPR and Cas6-based crRNA processing activity due to functional re-specialisation. Although the *M. sp*. JS623 type IV-B operon contains both Csf3 (Cas5) and the putative large subunit Csf1, we did not observe corresponding densities within our cryo-EM structure. In type I CRISPR systems, Cas5 binds the 5’ crRNA handle with high affinity and sequence specificity, nucleating complex assembly^3,29^. The lack of the Cas5-like Csf3 subunit within our complex may explain the heterogeneous assembly of type IV-B Csf complexes around non-specific RNA (**Supplementary Figure 2**). The ability of the Csf complex to assemble on non-specific RNAs suggests that type IV-B systems may have been functionally repurposed for a yet to be identified role.

Since all type IV systems identified lack adaptation subunits and almost all (97.8%) type IV-B operons identified lack any CRISPR array, it is likely they do not participate in selective pre-spacer acquisition or adaptive immunity^1,23^. Instead, they may have been co-opted for an orthogonal function. While there is a precedent for the hijacking and repurposing of CRISPR systems for non-defence functions^30^, the role of type IV-B systems remains a mystery. A particularly tantalising hypothesis is that type IV-B Csf complexes assemble on small RNAs, acting as non-specific RNA-sponges, and enabling IV-B-encoding megaplasmids to evade targeting by host cell RNA guided defences. Future experiments are essential to reveal the biological functions of type IV systems. Recent classifications have indicated that although type IV-B systems are highly diverse, they are almost always associated with an adenosine 5’-phosphosulfate reductase-family gene *cysH*^1,21,23^. Thus, understanding the interplay between *cysH* and the Csf complex may be the key to deciphering the enigmatic role of type IV-B CRISPR systems.

## Methods

### Expression and purification of the M. sp JS623 Csf complex

*E.coli* BL21 (DE3) cells containing the pCDF-Duet1-Csf1-Csf4-Strep-Csf2-Csf3(MCS1)-Cas6array(MCS2) expression vector were inoculated in 6 X 0.5 L lysogeny broth (LB) and grown at 37°C with 200 rpm shaking. Cells were grown to an optical density (O.D. 600 nm) between 0.6-0.7 then cold shocked on ice for 30-60 min. Recombinant protein expression was induced with 0.8 mM IPTG (isopropyl β-D-1-thiogalactopyranoside). After induction, cells were grown at 16°C for 18-24 hours and pelleted via centrifugation. Pelleted cells were resuspended in 20-30 mL Buffer W (100 mM Tris, pH 8.0; 150 mM NaCl; 2 *μ*M ZnSO_4_). Protease inhibitors were added to the following final concentrations: 10 *μ*g/mL leupeptin, 2 *μ*g/mL aprotinin, and 170 *μ*g/mL phenylmethylsulfonyl fluoride (PMSF). Cells were lysed by sonication and lysate was clarified by centrifugation. The soluble fraction was applied to a StrepTrap HP 5 mL column (GE Healthcare) and the bound protein was eluted with Buffer E (Buffer W + 5 mL desthiobiotin). The RNP complex was further purified with a Superose6 Increase 10/300 GL column (GE Healthcare), eluting into SEC Buffer (50 mM HEPES, pH 8.0; 150 mM NaCl; 2 *μ*M ZnSO_4_). Nucleic acids were extracted with acid phenol:chloroform and subsequent ethanol precipitation. The resulting fraction was loaded on a 20% denaturing PAGE gel after which a band of approximately 55-60 nt (Supplementary figure 2A) was excised and purified from gel using the ZR small-RNA PAGE Recovery Kit (Zymo Research, USA).

### RNA sequencing and analysis

Nucleic acids that co-purified with type IV-B complex were extracted with acid phenol:chloroform and subsequent ethanol precipitation. The resulting fraction was loaded on a 20% denaturing PAGE gel after which a band of approximately 55-60 nt (Supplementary figure 2A) was excised and purified from gel using the ZR small-RNA PAGE Recovery Kit (Zymo Research, USA). Small RNAs were prepared by GenXPro (GenXPro GmbH, Germany) using the TrueQuant smallRNA Seq kit according manufacturer’s instructions and were sequenced on a HiSeq2000 (Illumina, USA). After quality control filtering and adapter trimming using Cutadapt^31^, the reads were mapped on the expression plasmid and the *E.coli* BL21 (DE3) genome (Genbank accession CP001509) with Bowtie2^32^. Visualisation of the mapping results and further downstream analyses were performed using Geneious Prime 2020.10.2 (https://www.geneious.com).

### Cryo-EM data acquisition and processing

C-flat holy carbon grids (CF-4/2, Protochips Inc.) were glow-discharged for 30 seconds using a Gatan Solarus plasma cleaner. 2.5 μl of Type IV complex (~0.3 mg/ml) was applied onto grids, blotted for 2.5 seconds with a blotting force of 1 and rapidly plunged into liquid ethane using a FEI Vitrobot MarkIV operated at 4 °C and 100% humidity. Data were acquired using a FEI Titan Krios transmission electron microscope (Sauer Structural Biology Laboratory, University of Texas at Austin) operating at 300 keV at a nominal magnification of ×22,500 (1.1 Å pixel size) with defocus ranging from −1.5 to −3.0 μm. The data were collected using a total exposure of 6 s fractionated into 20 frames (300 ms per frame) with a dose rate of ~8 electrons per pixel per second and a total exposure dose of ~40 e^−^Å^−2^. Three datasets were automatically recorded on a Gatan K2 Summit direct electron detector operated in counting mode using the MSI-Template application within the automated macromolecular microscopy software LEGINON^33^.

All image pre-processing was performed in Appion^34^. Individual movie frames were aligned and averaged using ‘MotionCor2’ drift-correction software^35^. The contrast transfer function (CTF) of each micrograph was estimated using CTFFIND4^36^. Particles were picked with a template-based particle picker using a reference-free 2D class average from a small subset of manually picked particles as templates. Selected particles were extracted from micrographs using particle extraction within Relion^37^ and the EMAN2 coordinates exported from Appion. After multiple rounds of 2D classification in Relion to remove junk particles, 824,421 particles were left and imported into cryoSPARC^38^ for further processing. After multiple rounds of 2D classification and heterogeneous refinement, a final reconstruction containing 296,319 particles was determined to a global resolution of 3.9 Å (based on the gold standard 0.143 FSC criterion using two independent half-maps) using local refinement (implementing nonuniform refinement) with a mask corresponding to the entire complex.

### Csf model building, refinement and structural analysis

An atomic model for the Csf complex was built de novo in Coot^39^, and subjected to multiple iterative rounds of molecular dynamics - flexbile fitting in Namdinator^40^ and real-space refinement in Phenix^41^. The refined Csf complex model was validated using MolProbity^42^ as implemented in Phenix. Protein sequence conservation analysis was performed using online ConSurf^43^ server, with multiple sequence alignment (MSA) generated using the top 100 result from a BLAST search against Csf sequences. Output from the MSA was used to generate a sequence logo using the WebLogo server^44^. Maps and models were visualized using ChimeraX^45^ and the electrostatic surfaces were determined using the APBS plugin^46^ within PyMol. Root-mean-square deviation (r.m.s.d.) values between equivalent atoms in Csf2 and type III-A Csm3, and between type IV-B RNA and III-A crRNA were calculated using ChimeraX and PyMol.

## Acknowledgements

We thank members of the Staals, Jackson and Taylor labs for helpful discussions. This work was supported in part by Welch Foundation grant F-1938 (to D.W.T.), Army Research Office Grant W911NF-15-1-0120 (to D.W.T.), and a Robert J. Kleberg, Jr. and Helen C. Kleberg Foundation Medical Research Award (to D.W.T.). D.W.T is a CPRIT Scholar supported by the Cancer Prevention and Research Institute of Texas (RR160088) and an Army Young Investigator supported by the Army Research Office (W911NF-19-1-0021). Research in the Jackson Lab is supported by Utah State University New Faculty Start-up funding from the Department of Chemistry and Biochemistry, the Research and Graduate Studies Office, and the College of Science as well as the National Institutes of Health (R35GM138080). Data was collected at the Sauer Structural Biology Lab at UT Austin.

## Author contributions

H.N.T. performed purification of complexes. Y.Z. and J.P.K.B. collected and processed cryo-EM data and performed model-building and refinement. J.S. performed the RNA-seq experiments. All authors interpreted the results and wrote the manuscript. R.H.J.S., R.N.J., and D.W.T. conceived the experiments, supervised the research, and secured funding for the project.

## Competing interests

D.W.T. is an inventor on a CRISPR-related patent. All other authors declare no competing interests.

**Correspondence and requests for materials should be addressed to R.H.J.S., R.N.J., or D.W.T**.

## Data availability

The data that support the findings of this study are available from the corresponding authors upon request. The cryo-EM structure and associated atomic coordinates have been deposited in the Electron Microscopy Data Bank and the Protein Data Bank with accession codes EMD-YYYYY and ZZZZ, respectively.

## Supplementary Information

**Supplementary Figure 1.**
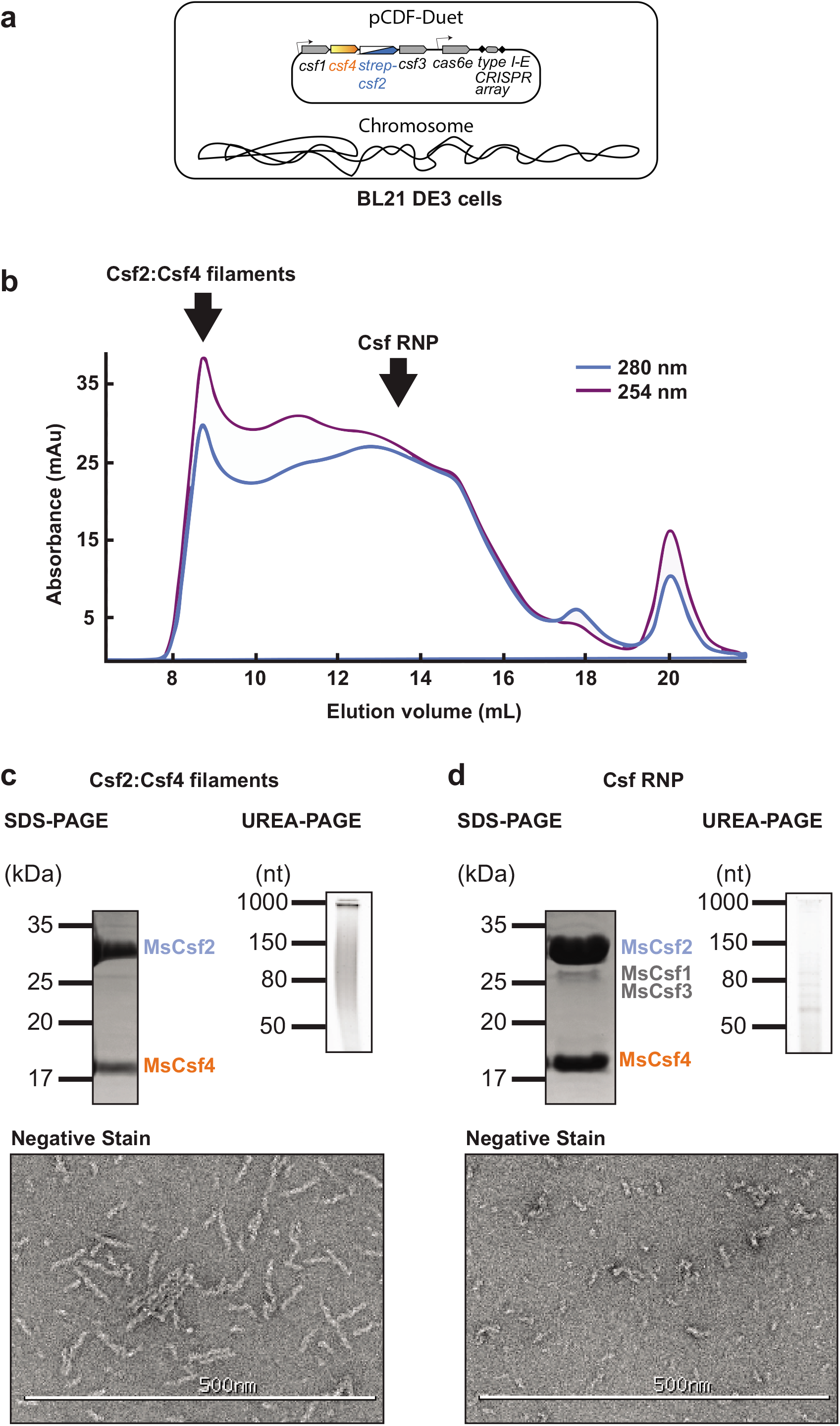
Purification of the *M. sp JS623* RNP complex by affinity chromatography (N-Strep-MsCsf2) and size exclusion chromatography. **a**, Diagram of plasmid used to express the type IV-B complex in BL21 DE3 cells. **b**, SEC chromatogram highlighting peaks corresponding to Csf2:Csf4 filaments and the RNP complex. **c**, SDS-PAGE, UREA-PAGE, and negative stain data indicating the presence of Csf2:Csf4 bound to long RNAs to create filamentous structures. **d**, SDS-PAGE, UREA-PAGE, and negative stain data indicating the presence of Csf1, Csf2, Csf3, and Csf4 bound to short, distinct RNAs to create an RNP complex.

**Supplementary Figure 2.**
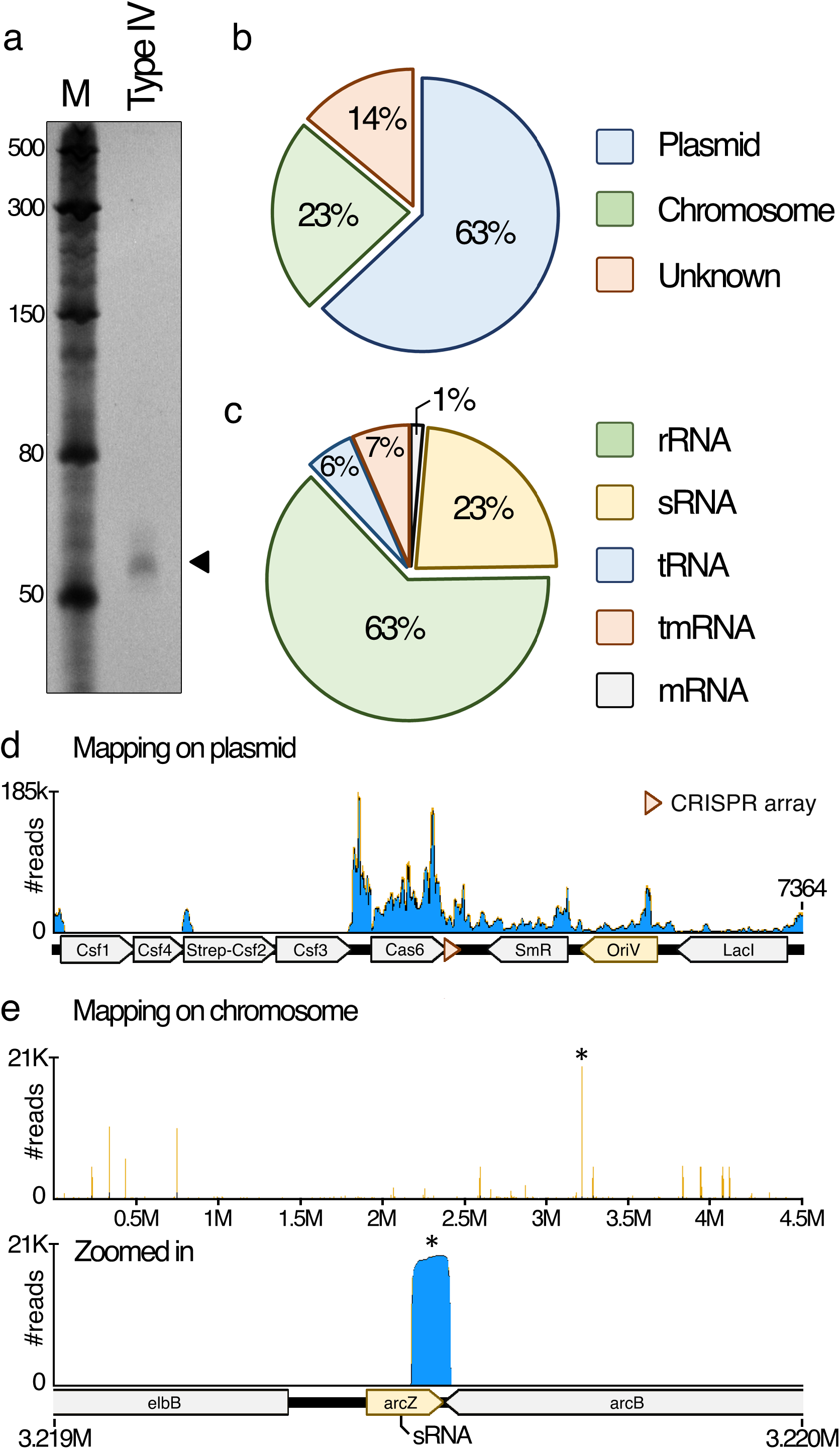
RNA sequencing on co-purifying nucleotides with type IV complex. **a**, UREA-PAGE gel showing nucleic acids co-purifying with. The triangle indicates the ~60nt band purified from isolated type IV-B complexes that was used for RNA sequencing analysis. **b**, Percentage of reads mapping to either to expression plasmid, chromosome, or of unknown origin. **c**, Origins of reads mapped on the chromosome, based on the type of RNA. Classes of RNA found to be associated with the type IV complex: Ribosomal RNA (rRNA), small regulatory RNA (sRNA), transfer RNA (tRNA), transfer-messenger RNA (tmRNA), messenger RNA (mRNA). **d**, Distribution of reads mapping on the expression plasmid. **e**, Top panel -Distribution of reads mapping on the chromosome. Highlighted by the asterix (*) is the location on the chromosome with the highest abundance of reads (~21k). Lower panel – A zoomed in representation of the area with the highest abundance of reads, *arcZ*, a known short regulatory RNA (sRNA).

**Supplementary Figure 3.**
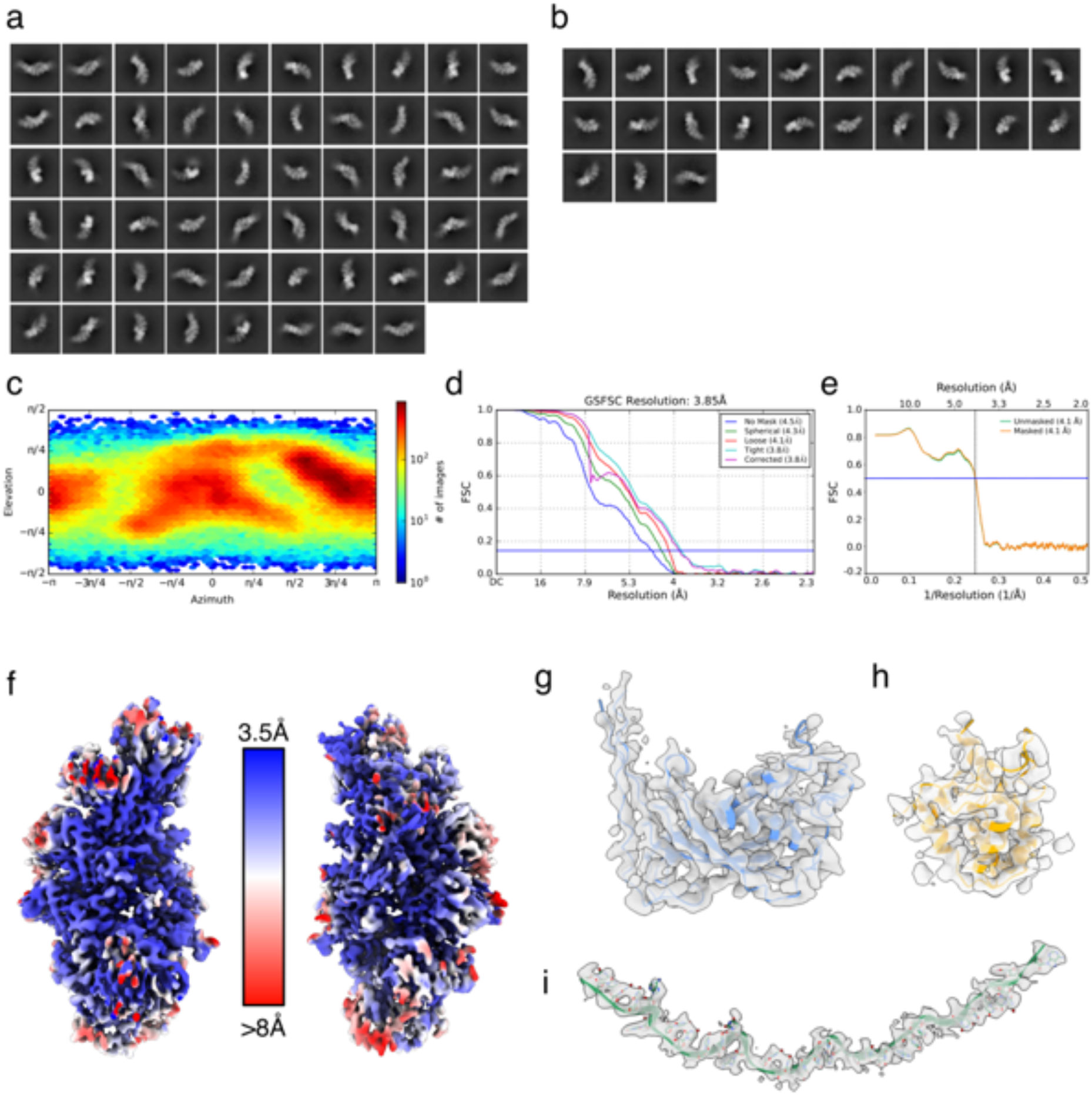
Cryo-EM analysis of Csf. **a**, 2D class averages of autopicked particles. **b**, 2D class averages of discrete Csf oligomers used for 3D reconstruction. **c**, Euler angular distribution of particles contributing to final 3D reconstruction. **d**, Fourier shell correlation (FSC) of final 3D reconstruction, with a global resolution of 3.9 Å. **e**, Map-to-model FSC, with a resolution of 4.1Å at FSC threshold of 0.5. **f**, Map of Csf complex colored by local resolution. **g – i**: Representative atomic models and corresponding cryo-EM densities for Csf2 (**g**), Csf4 (**h**) and RNA (**i**).

**Supplementary Figure 4.**
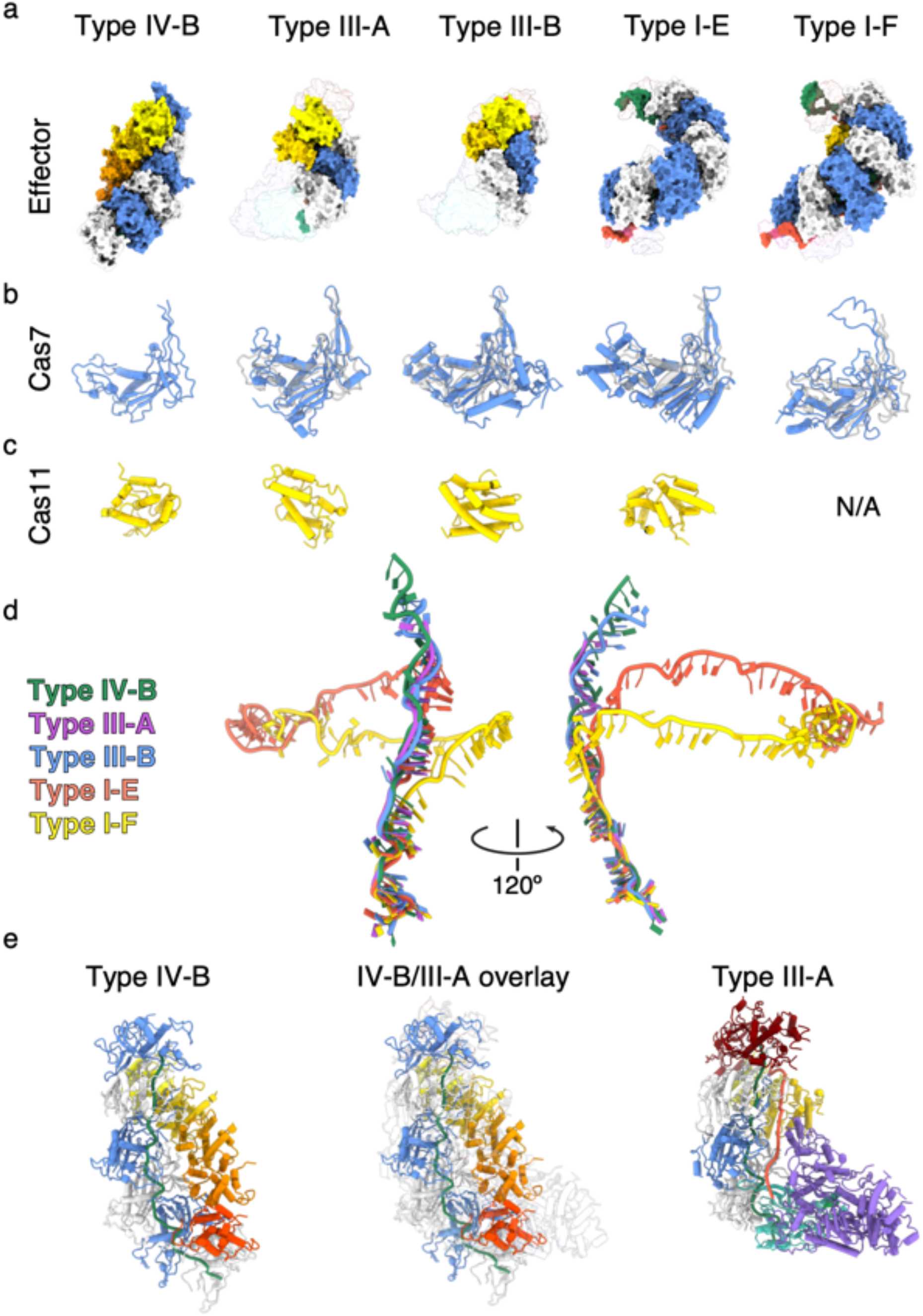
Comparison to Cascade complexes. **a**, Comparison of Type IV-B Csf with other effector/Cascade complexes. crRNA within Type III-A (PDB 607i) & III-B (PDB 3×1l) complexes were aligned to Csf RNA (RMSD 5.9 Å and 8.8 Å, respectively). Due to the highly curved nature of type I-E (5h9f) and I-F (6b44) crRNA, it was not possible to perform such alignment to IV-B. Instead, individual backbone subunits were aligned to corresponding Csf2 subunits, thus aligning the top of I-E or I-F with the top of IV-B (RMSD ~24.4 Å and 25.8 Å, respectively). In all complexes, non-Cas7/Cas11 subunits are shown as transparent surfaces. All Cas7/Cas11 and RNA, crRNA and TS are colored as in **Figures 1 & 2**, with the addition of the TS in light red. **b**, Alignment of Cas7 with Csf2. Csf2 is shown as grey, transparent cartoon. RMSD is typically ~20 Å – 25 Å, although they clearly align well by eye. The high RMSD is likely due to presence of additional residues not present in Csf2. **c**, Cas11 subunits. All Cas11 subunits are helical bundles that resemble each other. However, due to diverse Cas11 sequences these subunits align poorly (RMSD 16 Å – 18 Å)**. d**, Alignment of (cr)RNA from available CRISPR effector complexes. Type IV-B RNA aligns more closely to type III-A and -B crRNAs, consistent with the proposed evolutionary lineage of type IV CRISPR systems emerging from a type III-like ancestor. **e**, Overlay of type IV-B and type III-A CRISPR complexes based on RNA alignment. Type III-A displays the strongest structural homology between Cas7 subunits.

**Supplementary Figure 5.**
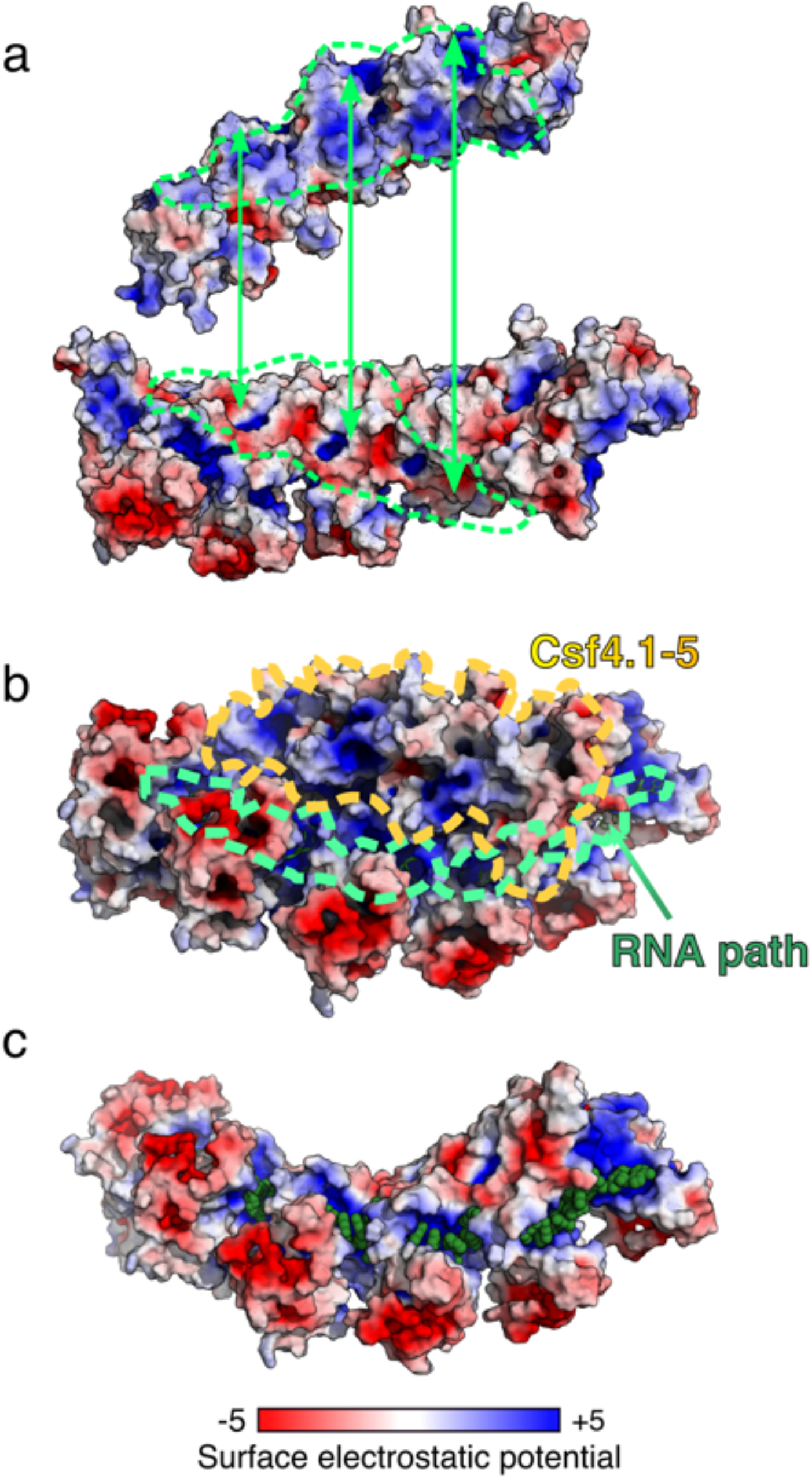
Surface electrostatics of Csf complex. **a & b**, Csf4 and Csf2 filaments. Green dashed outlines denote complementary surfaces. **b**, Surface electrostatics of the Csf complex. Csf4 subunits and RNA path are outlined. RNA is shown as green cartoon, but it is almost completely occluded by Cas7 subunits. The overall path of the RNA bound within the Cas7 filament is outlined by green dashed lines. **c**, Csf2 filament with RNA contacts (green spheres) shown. Csf4 minor filament is removed for clarity. RNA is bound by a contiguous positively-charged surface. The high electrostatic contribution to RNA binding by Csf2 is typical of non-specific RNA-binding proteins^47^.

**Supplementary Figure 6.**
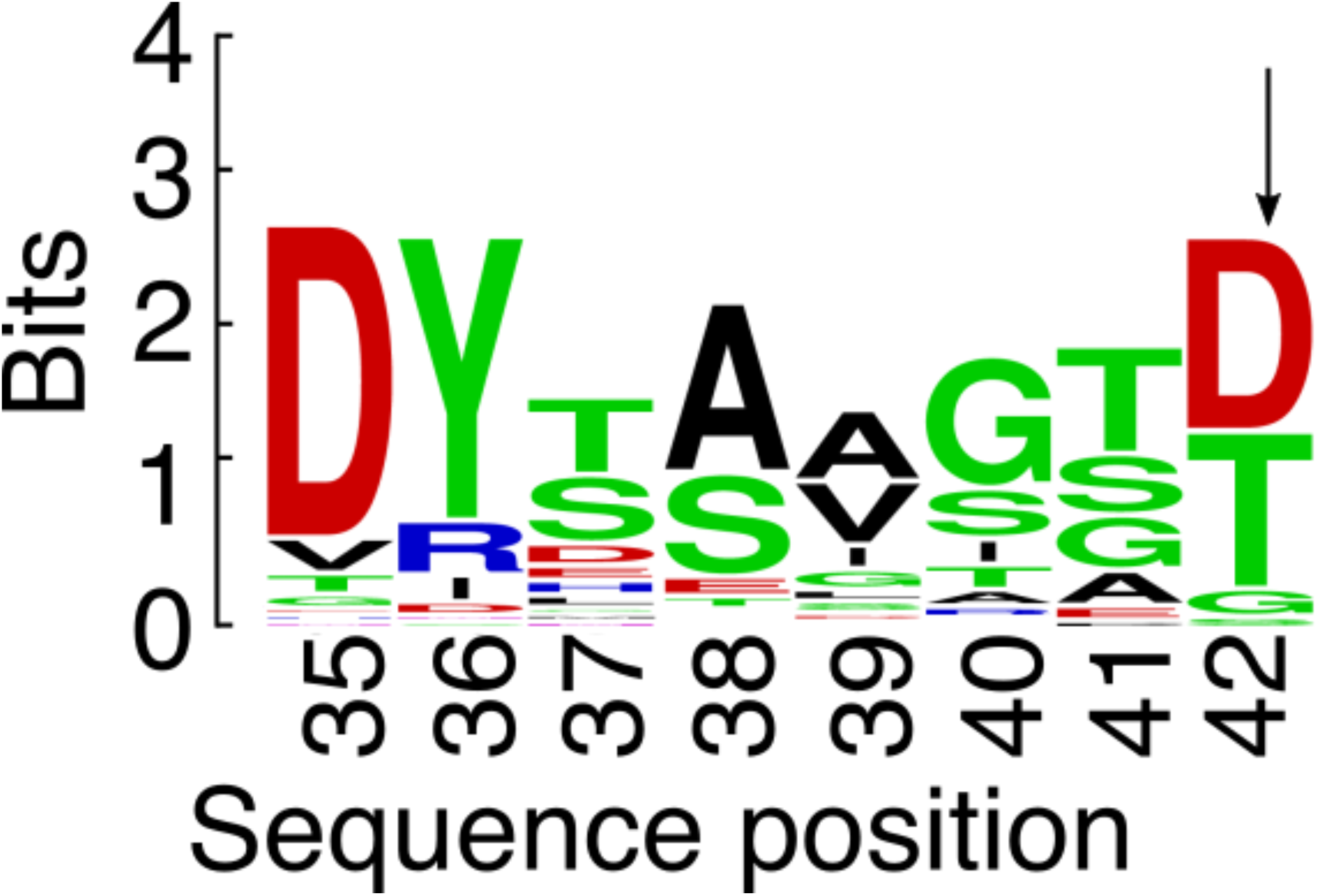
Weblogo of cleavage loop, with candidate catalytic residue (D42) denoted by arrow. Multiple sequence alignment (MSA) generated using the top 100 result from a BLAST search against the Csf2 sequence from *M. sp JS623*.. Output from the MSA was used to generate a sequence logo using the WebLogo server^44^ and conservation score used in Figure 2.

**Supplementary Table 1.** Cryo-EM data collection and processing parameters.

| **Data collection and processing** |  |
| --- | --- |
| Magnification | 22,500x |
| Voltage (kV) | 300 |
| Electron exposure (e<sup>-</sup>/Å) | 40 |
| Defocus range (μM) | -1.5 to -3.0 |
| Symmetry imposed | C1 |
| Initial particle images | 824,221 |
| Final particle images | 296,319 |
| Map resolution (Å) | 3.9 |
| FSC threshold | 0.143 |
| Map resolution range (Å) | 3.5 to > 8 |
| **Refinement** |  |
| Initial model used (PBB code) | N/A |
| Model resolution (Å) | 4.1 |
| FSC threshold | 0.5 |
| Map sharpening *B* factor (Å<sup>2</sup>) | 130.2 |
| **Model composition** |  |
| Nonhydrogen atoms | 16636 |
| Residues (Protein/RNA) | 2083/31 |
| ***B* factors (Å<sup>2</sup>) (min/max/mean)** |  |
| Protein | 19.75/206.87/81.59 |
| RNA | 34.71/125.88/56.91 |
| **r.m.s. deviations** |  |
| Bond lengths (Å) | 0.005 |
| Bond angles (°) | 1.122 |
| **Validation** |  |
| MolProbity score | 1.9 |
| Clashscore | 4.72 |
| Poor rotamers (%) | 0 |
| **Ramachandran plot** |  |
| Favoured (%) | 84.99 |
| Allowed (%) | 15.01 |
| Disallowed (%) | 0 |

